# Selection at the pathway level drives the evolution of gene-specific transcriptional noise

**DOI:** 10.1101/104216

**Authors:** Gustavo Valadares Barroso, Natasa Puzovic, Julien Y Dutheil

**Affiliations:** Max Planck Institute for Evolutionary Biology. Department of Evolutionary Genetics. August-Thienemann-Straße 2 24306 Plön – GERMANY; ISEM – Institut des Sciences de l’Évolution. UMR 5554, Université de Montpellier, Place Eugène Bataillon 34095 Montpellier cedex 05 – FRANCE

## Abstract

Biochemical reactions within individual cells result from the interactions of molecules, typically in small numbers. Consequently, the inherent stochasticity of binding and diffusion processes generate noise along the cascade that leads to the synthesis of a protein from its encoding gene. As a result, isogenic cell populations display phenotypic variability even in homogeneous environments. The extent and consequences of this stochastic gene expression have only recently been assessed on a genome-wide scale, in particular owing to the advent of single cell transcriptomics. However, the evolutionary forces shaping this stochasticity have yet to be unraveled. We take advantage of two recently published data sets of the single-cell transcriptome of the domestic mouse *Mus musculus* in order to characterize the effect of natural selection on gene-specific transcriptional stochasticity. We show that noise levels in the mRNA distributions (*a.k.a.* transcriptional noise) significantly correlate with three-dimensional nuclear domain organization, evolutionary constraint on the encoded protein and gene age. The position of the encoded protein in biological pathways, however, is the main factor that explains observed levels of transcriptional noise, in agreement with models of noise propagation within gene networks. Because transcriptional noise is under widespread selection, we argue that it constitutes an important component of the phenotype and that variance of expression is a potential target of adaptation. Stochastic gene expression should therefore be considered together with mean expression level in functional and evolutionary studies of gene expression.

## Introduction

Isogenic cell populations display phenotypic variability even in homogeneous environments (Spudich and Koshland 1976). This observation challenged the clockwork view of the intra-cellular molecular machinery and led to the recognition of the stochastic nature of gene expression. Since biochemical reactions result from the interactions of individual molecules in small numbers (Gillesple 1977), the inherent stochasticity of binding and diffusion processes generates noise along the biochemical cascade leading to the synthesis of a protein from its encoding gene (**Figure 1**). The study of stochastic gene expression (SGE) classically recognizes two sources of expression noise. Following the definition introduced by Elowitz et al (Elowitz et al. 2002), extrinsic noise results from variation in concentration, state and location of shared key molecules involved in the reaction cascade from transcription initiation to protein folding. This is because molecules that are shared among genes, such as ribosomes and RNA polymerases, are typically present in low copy numbers relative to the number of genes actively transcribed (Shahrezaei and Swain 2008). Extrinsic factors also include physical properties of the cell such as size and growth rate, likely to impact the diffusion process of all molecular players. Extrinsic factors therefore affect every gene in a cell equally. Conversely, intrinsic factors generate noise in a gene-specific manner. They involve, for example, the strength of cis-regulatory elements (Suter et al. 2011) as well as the stability of the mRNA molecules that are transcribed (Mcadams and Arkin 1997; Thattai and Oudenaarden 2001). Every gene is affected by both sources of stochasticity and the relative importance of each has been discussed in the literature (Becskei et al. 2005; Raj and Oudenaarden 2008). Shahrezaei and Swain (Shahrezaei and Swain 2008) proposed a more general, systemic and explicit definition for any organization level, where intrinsic stochasticity is “generated by the dynamics of the system from the random timing of individual reactions” and extrinsic stochasticity is “generated by the system interacting with other stochastic systems in the cell or its environment”. This generic definition therefore includes Raser and O’Shea’s (Raser and O’Shea 2005) suggestion to further distinguish extrinsic noise occurring “within pathways” and “between pathways”. Other organization levels of gene expression are also likely to affect expression noise, such as chromatin structure (Blake et al. 2003; Hebenstreit 2013), and three-dimensional genome organization (Pombo and Dillon 2015).

**Figure 1:**
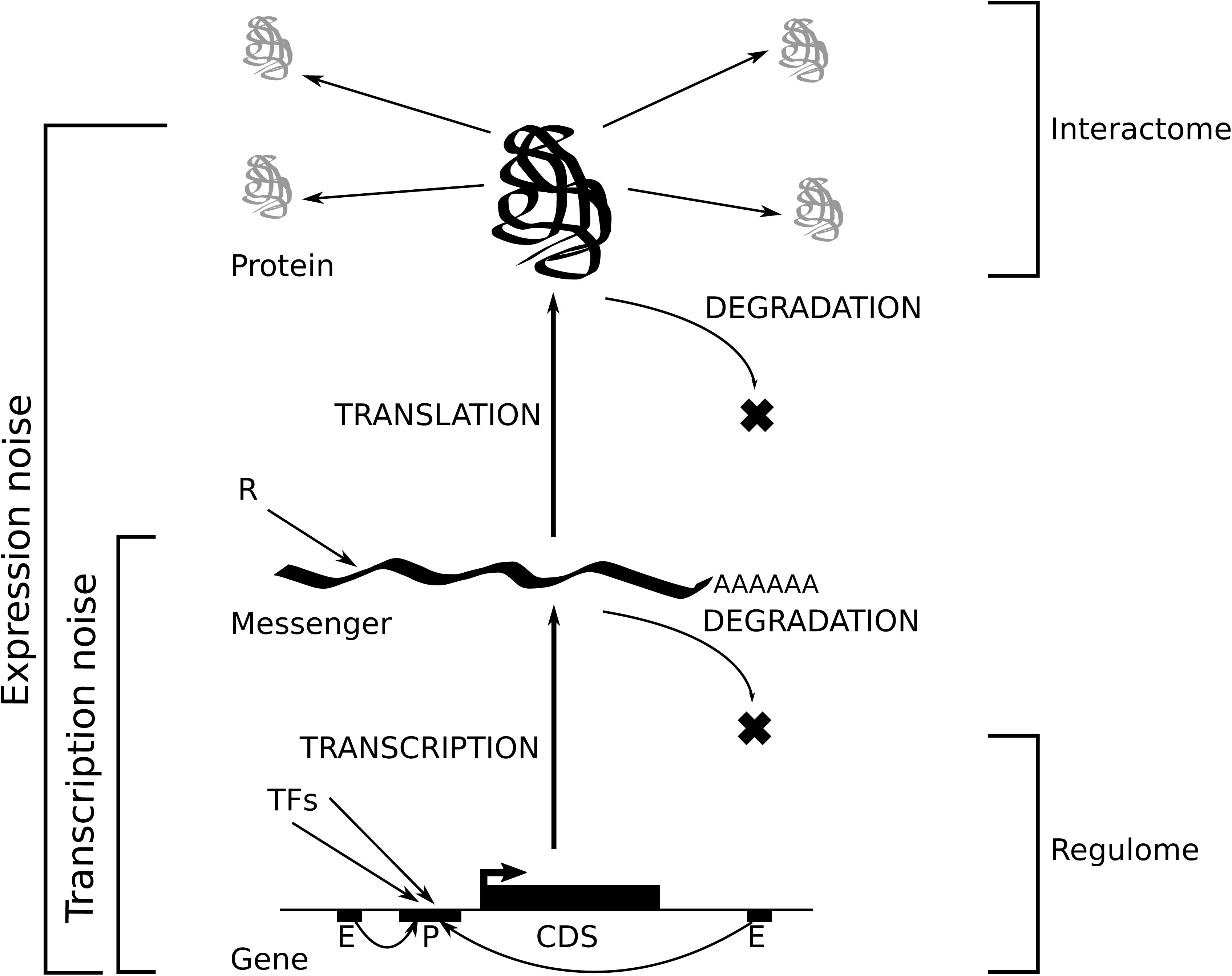
A systemic view of gene expression.

Pioneering work by Fraser et al (Fraser et al. 2004) has shown that SGE is an evolvable trait which is subject to natural selection. First, genes involved in core functions of the cell are expected to behave more deterministically (Barkai and Leibler 1999) because temporal oscillations in the concentration of their encoded proteins are likely to have a deleterious effect. Second, genes involved in immune response (Arkin et al. 1998; Norman et al. 2015) and response to environmental conditions can benefit from being unpredictably expressed in the context of selection for bet-hedging (Thattai and Oudenaarden 2004). As the relation between fitness and stochasticity depends on the function of the underlying gene, selection on SGE is expected to act mostly at the intrinsic level (Newman et al. 2006; Lehner 2008; Wang and Zhang 2011). The molecular mechanisms by which natural selection operates to regulate expression noise, however, remain to be elucidated.

Due to methodological limitations, seminal studies on SGE (both at the mRNA and protein levels) have focused on only a handful of genes (Elowitz et al. 2002; Ozbudak et al. 2002; Chubb et al. 2006). The canonical approach consists in selecting genes of interest and recording the change of their noise levels in a population of clonal cells as a function of either (1) the concentration of the molecule that allosterically controls affinity of the transcription factor to the promoter region of the gene (Blake et al. 2003; Bar-even et al. 2006) or (2) mutations artificially imposed in regulatory sequences (Ozbudak et al. 2002). In parallel with theoretical work (Kepler and Elston 2001; Batada and Hurst 2007; Kaufmann and van Oudenaarden 2007; Sánchez and Kondev 2008), these pioneering studies have provided the basis of our current understanding of the proximate molecular mechanisms behind SGE, namely complex regulation by transcription factors, architecture of the upstream region (including the presence of TATA box) and gene orientation (Wang et al. 2011), translation efficiency and mRNA / protein stability (Eldar and Elowitz 2010), properties of the protein-protein interaction network (Li et al. 2010). Measurements at the genome scale coupled with rigourous statistical analyses are however needed in order to go beyond gene idiosyncrasies and particular histories, and test hypotheses about the evolutionary forces shaping SGE (Sauer et al. 2007).

The recent advent of single-cell RNA sequencing makes it possible to sequence the transcriptome of each individual cell in a collection of clones, and to observe the variation of gene-specific mRNA quantities across cells. This provides a genome-wide assessment of transcriptional noise. While not accounting for putative noise resulting from the process of translation of mRNAs into proteins, transcriptional noise accounts for noise generated by both synthesis and degradation of mRNA molecules (**Figure 1**). Previous studies, however, have shown that transcription is a limiting step in gene expression, and that transcriptional noise is therefore a good proxy for expression noise (Newman et al. 2006; Taniguchi et al. 2011). Here, we used publicly available single-cell transcriptomics data sets to quantify gene-specific transcriptional noise and relate it to other genomic factors, including protein conservation and position in the interaction network, in order to uncover the molecular basis of selection on stochastic gene expression.

## Results

### A new measure of noise to study genome-wide patterns of stochastic gene expression

We used the dataset generated by Sasagawa et al (2013), which quantifies gene-specific amounts of mRNA as fragments per kilobase of transcripts per million mapped fragments (FPKM) values for each gene and each individual cell. Among these, we selected all genes in a subset containing 20 embryonic stem cells in G1 phase in order to avoid recording variance that is due to different cell types or cell-cycle phases. The Quartz-Seq sequencing protocol captures every poly-A RNA present in the cell at one specific moment, allowing to assess transcriptional noise. Following Shalek et al (2014) we first filtered out genes that were not appreciably expressed in order to reduce the contribution of technical noise to the total noise. For each gene we further calculated the mean *μ* in FPKM units and variance *σ* ^2^ in FPKM ^2^ units, as well as two previously published measures of stochasticity: the *Fano factor*, usually referred to as the bursty parameter, defined as *σ* ^2^/ *μ* and *Noise*, defined as the coefficient of variation squared (*σ* ^2^/ *μ* ^2^). Both the variance and *Fano factor* are monotonically increasing functions of the mean (**Figure 2A**). *Noise* is inversely proportional to mean expression (**Figure 2A**), in agreement with previous observations at the protein level (Bar-even et al. 2006; Taniguchi et al. 2011). While this negative correlation was theoretically predicted (Tao et al. 2007), it may confound the analyses of transcriptional noise at the genome level, because mean gene expression is under specific selective pressure (Pál et al. 2001). In order to disentangle these effects, we developed a new quantitative measure of noise, independent of the mean expression level of each gene. To achieve this we performed polynomial regressions in the log-space plot of variance *versus* mean. We defined F* as 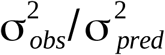 (see Material and Methods) that is, the ratio of the observed variance over the variance component predicted by the mean expression level. We selected the simplest model for which no correlation between F* and mean expression was observed, and found that a degree 3 polynomial model was sufficient to remove further correlation (Kendall’s tau = -0.0037, p-value = 0.5217, **Figure 2A**). Genes with F* < 1 have a variance lower than expected according to their mean expression whereas genes with F* > 1 behave the opposite way **(Figure 2B**). This approach fulfills the same goal as the running median approach of Newman et. al (Newman et al. 2006), whilst it includes the effect of mean expression directly into the measure of stochasticity instead of correcting a posteriori a dependent measure (in that case, the Fano factor). We therefore use F* as a measure of SGE throughout this study.

**Figure 2:**
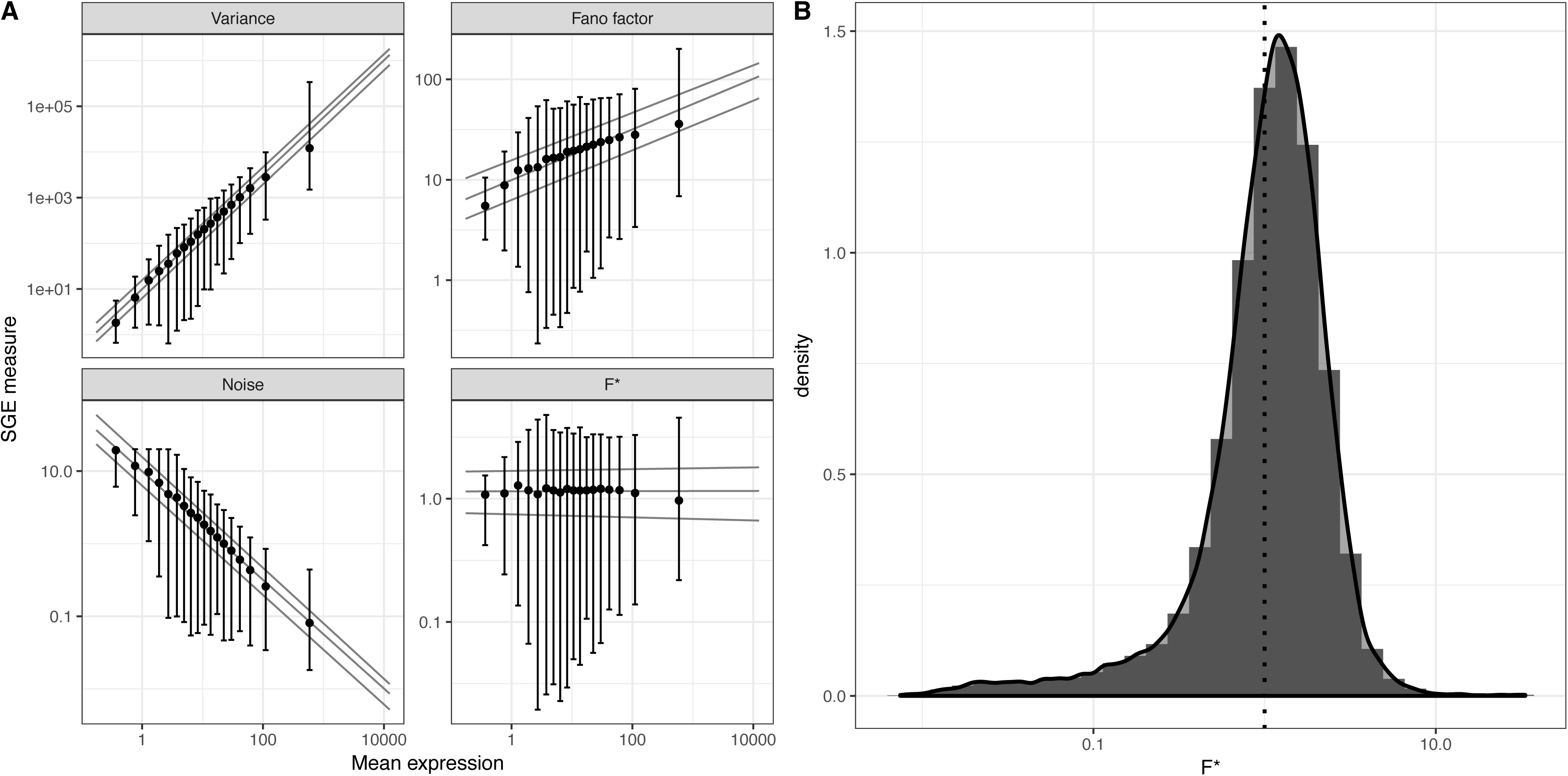
Transcriptional noise and mean gene expression. A) Measures of noise plotted against the mean gene expression for each gene, in logarithmic scales: Variance, Fano factor (variance / mean), noise (square of the coefficient of variation, variance / mean^2) and F* (this study). Lines represent quantile regression fits (median, first and third quartiles). Point and bars represent median, first and third quartiles for each category of mean expression obtained by discretization of the x axis. B) Distribution of F* over all genes in this study. Vertical line corresponds to F* = 1.

### Stochastic gene expression correlates with the three-dimensional structure of the genome

We first sought to investigate whether genome organization significantly impacts the patterns of stochastic gene expression. We assessed whether genes in proximity along chromosomes display more similar amount of transcriptional noise than distant genes. We tested this hypothesis by computing the primary distance on the genome between each pair of genes, that is, the number of base pairs separating them on the chromosome, as well as the relative difference in their transcriptional noise (see Methods). We found no significant association between the two distances (Mantel tests, each chromosome tested independently). Contiguous genes in one dimension, however, have significantly more similar transcriptional noise that non-contiguous genes (permutation test, p-value < 1e-4, **Figure S1**). Using Hi-C data from mouse embryonic cells (Dixon et al. 2012), we report that genes in contact in three-dimensions have significantly more similar transcriptional noise than genes not in contact (permutation test, p-value < 1e-3, **Figure S1**). Most contiguous genes in one-dimension also appear to be close in three-dimensions and the effect of 3D contact is stronger than that of 1D contact. These results therefore suggest that the three-dimensional structure of the genome has a stronger impact on stochastic gene expression than the position of the genes along the chromosomes. We further note that while highly significant, the size of this effect is small, with a difference in relative expression of -1.10% (**Figure S1**).

### Transcription factors binding and histone methylation impact stochastic gene expression

The binding of transcription factors (TF) to promoter constitutes one notable source of transcriptional noise (**Figure 1**) (Blake et al. 2003; Newman et al. 2006). In eukaryotes, the accessibility of promoters is determined by the chromatin state, which is itself controlled by histone methylation. We assessed the extent to which transcriptional noise is linked to particular TFs and histone marks by using data from the Ensembl regulatory build (Zerbino et al. 2015), which provides data from experimental evidence of TF binding and methylation sites along the genome. First we contrasted the F* values of genes with binding evidence for each annotated TF independently. Among 13 TF represented by at least 5 genes in our data set, we found that 4 of them significantly influence F* after adjusting for a global false discovery rate of 5%: the transcription repressor CTFC (adjusted p-value = 0.0321), the transcription factor CP2-like 1 (Tcfcp2l1, adjusted p-value = 0.0087), the X-Linked Zinc Finger Protein (Zfx, adjusted p-value = 0.0284) and the Myc transcription factor (MYC, ajusted p-value = 0.0104). Interestingly, association with each of these four TFs led to an increase in transcriptional noise. We also report a weak but significant positive correlation between the number of transcription factors associated with each gene and the amount of transcriptional noise (Kendall’s tau = 0.0238, p-value = 0.0007). This observation is consistent with the idea that noise generated by each TF is cumulative (Sharon et al. 2014). We then tested if particular histone marks are associated with transcriptional noise. Among five histone marks represented in our data set, three were found to be highly significantly associated to a higher transcriptional noise: H3K4me3 (adjusted p-value = 1.9981e-146), H3K4me2 (adjusted p-value = 5.4524e-121) and H3K27me3 (adjusted p-value = 5.2985e-34). Methylation on the fourth Lysine of histone H3 is associated with gene activation in humans, while tri-methylation on lysine 27 is usually associated with gene repression (Barski et al. 2007). These results suggest that both gene activation and silencing contribute to the stochasticity of gene expression, in agreement with the view that bursty transcription leads to increased noise (Blake et al. 2003; Newman et al. 2006; Batada and Hurst 2007).

### Low noise genes are enriched for housekeeping functions

We investigated the function of genes at both ends of the F* spectrum. We defined as candidate gene sets the top 10% least noisy or the top 10% most noisy genes in our data set, and tested for enrichment of GO terms and Reactome pathways (see Methods). It is expected that genes encoding proteins participating in housekeeping pathways are less noisy because fluctuations in concentration of their products might have stronger deleterious effects (Pedraza and van Oudenaarden 2005). On the other hand, stochastic gene expression could be selectively advantageous for genes involved in immune and stress response, as part of a bet-hedging strategy (eg Arkin et al. 1998; Shalek et al. 2013). GO terms enrichment test revealed significant categories enriched in the low noise gene set only: molecular functions “nucleic acid binding” and “structural constituent of ribosome”, the biological processes “nucleosome assembly”, “innate immune response in mucosa” and “translation”, as well as the cellular component “cytosolic large ribosomal subunit” (**Table 1**). All these terms but one relate to gene expression, in agreement with previously reported findings in yeast (Newman et al. 2006). We further find a total of 41 Reactome pathways significantly over-represented in the low-noise gene set (false discovery rate set to 1%). Interestingly, the top most significant pathways belong to modules related to translation (RNA processing, initiation of translation and ribosomal assembly), as well as several modules relating to gene expression, including chromatin regulation and mRNA splicing (**Figure 3**). Only one pathway was found to be enriched in the high noise set: TP53 regulation of transcription of cell cycle genes (p-value = 0.0079). This finding is interesting because TP53 is a central regulator of stress response in the cell (Hussain and Harris 2006). These results therefore corroborate previous findings that genes involved in stress response might be evolving under selection for high noise as part of a bet hedging strategy (Shalek et al. 2013; Viney and Reece 2013). The small amount of significantly enriched Reactome pathways by high noise genes can potentially be explained by the nature of the data set: as the original experiment was based on unstimulated cells, genes that directly benefit from high SGE might not be expressed in these experimental conditions.

**Figure 3:**
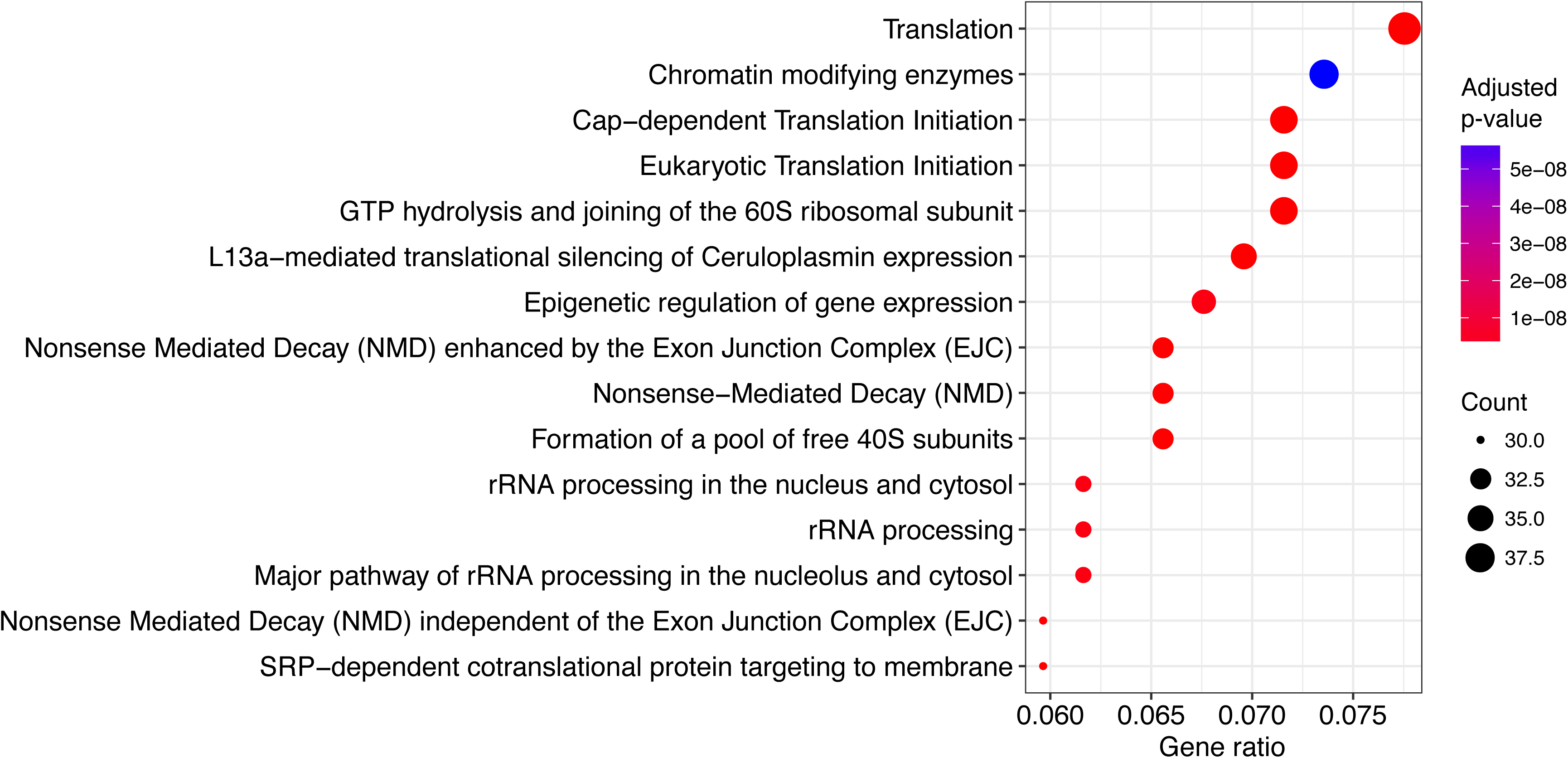
Enriched pathways in the low-noise gene set. Depicted pathways are the fifteen most significant in the 10% genes with lowest transcriptional noise.

**Table 1:**
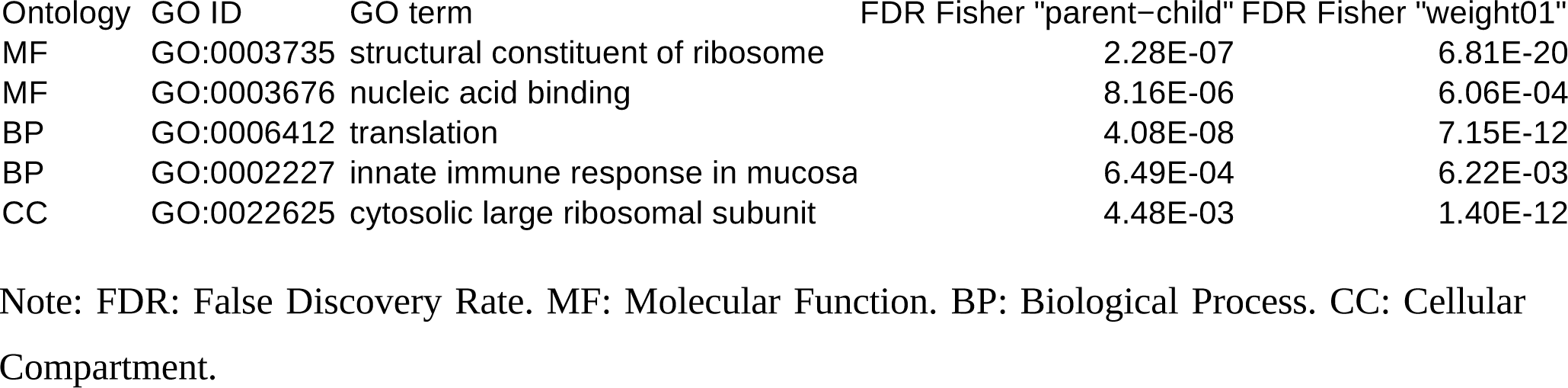
GO terms significantly enriched in the 10% genes with lowest transcriptional noise.

### Highly connected proteins are synthesized by low-noise genes

The structure of the interaction network of proteins inside the cell can greatly impact the evolutionary dynamics of genes (Jeong et al. 2000; Barabási and Oltvai 2004). Furthermore, the contribution of each constitutive node within a given network varies. This asymmetry is largely reflected in the power-law-like degree distribution that is observed in virtually all biological networks (Barabási and Albert 1999) with a few genes displaying many connections and a majority of genes displaying only a few. The individual characteristics of each node in a network can be characterized by various measures of centrality (Newman 2003). Following previous studies on protein evolutionary rate (Fraser et al. 2002; Hahn et al. 2004; Jovelin and Phillips 2009) and protein-protein interaction (PPI) networks (Li et al. 2010) we asked whether, at the gene level, there is a link between centrality of a protein and the amount of transcriptional noise. We study six centrality metrics measured on two types of network data: (i) pathway annotations from the Reactome database (Fabregat et al. 2016) and (ii) PPI data from the iRefIndex database. PPI data are typically more complete (5,553 genes with gene expression data) but do not provide functional evidence. The Reactome database is based on published functional evidence, but encompasses less genes (4,454 genes for which expression data is available). In addition, graph representing PPI network are not oriented while graph representing Pathway annotations are, implying that distinct statistics can be computed on both types of networks.

We first estimated the pleiotropy index of each gene by counting how many different pathways the corresponding proteins are involved in. We then computed centrality measures as averages over all pathways in which each gene is involved. These measures include (1) *node degree*, which corresponds to the number of other nodes a given node is directly connected with, (2) *hub score*, which estimates the extent to which a node links to other central nodes, (3) *authority score*, which estimates the importance of a node by assessing how many hubs link to it, (4) *transitivity*, or *clustering coefficient*, defined as the proportion of neighbors that also connect to each other, (5) *closeness*, a measure of the topological distance between a node and every other reachable node (the fewer edge hops it takes for a protein to reach every other protein in a network, the higher its closeness), and (6) *betweenness*, a measure of the frequency with which a protein belongs to the shortest path between every pairs of nodes.

We find that node degree, hub score, authority score and transitivity are all significantly negatively correlated with transcriptional noise on pathway-based networks: the more central a protein is, the less transcriptional noise it displays (**Figure 4A-D** and **Table 2**). We also observed that pleiotropy is negatively correlated with F* (Kendall’s tau = -0.0514, p-value = 8.31E-07, **Figure 4E**, **Table 2**), suggesting that a protein that potentially performs multiple functions at the same time needs to be less noisy. This effect is not an artifact of the fact that pleiotropic genes are themselves more central (e.g. correlation of pleiotropy and node degree: Kendall’s tau = 0.2215, p-value < 2.2E-16) or evolve more slowly (correlation of pleiotropy and Ka / Ks ratio: Kendall’s tau = -0.1060, p-value < 2.2E-16) since it is still significant after controlling for these variables (partial correlation of pleiotropy and F*, accounting for centrality measures and Ka / Ks: Kendall’s tau = -0.0254, p-value = 7.45E-06). Closeness and betweenness, on the other hand, show a negative correlation with F*, yet much less significant (Kendall’s tau = -0.0254, p-value = 0.0109 for closeness and tau = -0.0175, p-value = 0.0865 for betweenness, see **Figure 4FG** and **Table 2**). In modular networks (Hartwell et al. 1999) nodes that connect different modules are extremely important to the cell (Guimera and Amaral 2005) and show high betweenness scores. In yeast, high betweenness proteins tend to be older and more essential (Joy et al. 2005), an observation also supported by our data set (betweenness *vs* gene age, Kendall’s tau = 0.0619, p-value = 1.09E-07; betweenness *vs* Ka/Ks, Kendall’s tau = -0.0857, p-value = 3.83E-16). It has been argued, however, that in protein-protein interaction networks high betweenness proteins are less essential due to the lack of directed information flow, compared to, for instance, regulatory networks (Yu et al. 2007), a hypothesis which could explain the observed lack of correlation.

**Figure 4:**
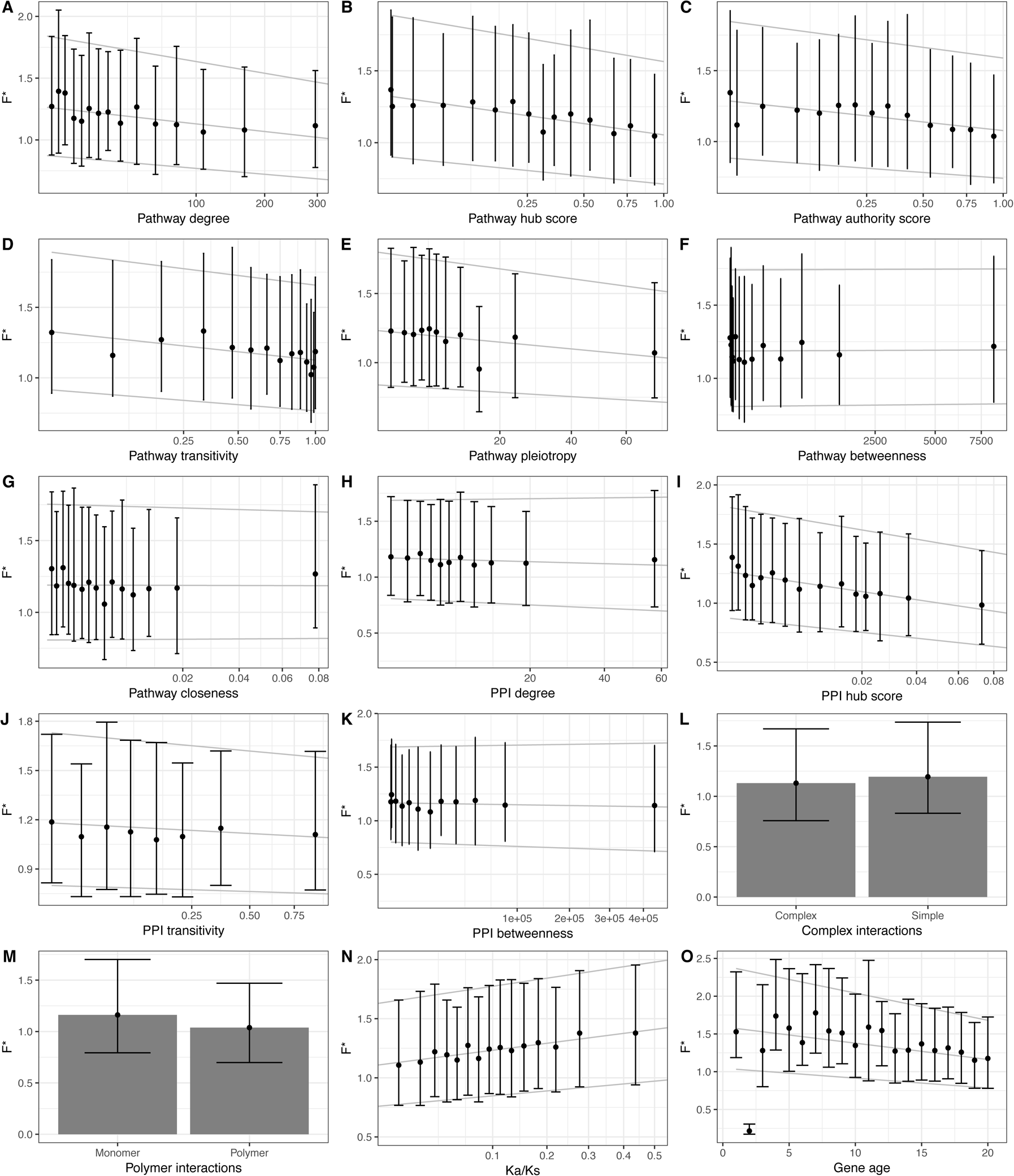
Factors driving stochastic gene expression. Correlation of F* and all tested network centrality measures, as well as protein conservation (Ka / Ks ratio) and gene age. Point and bars represent median, first and third quartiles for each category of mean expression obtained by discretization of the x axis, together with the quantile regression lines estimated on the full data set.

**Table 2:** Correlation of transcriptional noise with genes centrality measures and pleiotropy, as estimated from pathway annotations and protein-protein interactions networks.

| Data | Measure | Correlation with F* | p-value |
| --- | --- | --- | --- |
| Pathways | Degree | -0.0745 | 1.14E-13 *** |
|  | Hub score | -0.0808 | 6.61E-16 *** |
|  | Authority score | -0.0666 | 2.72E-11 *** |
|  | Clustering coefficient | -0.0794 | 4.55E-15 *** |
|  | Closeness | -0.0254 | 1.09E-02 * |
|  | Betweenness | -0.0175 | 8.65E-02 . |
|  | Pleiotropy | -0.0514 | 8.31E-07 *** |
|  | Size | -0.0514 | 3.91E-03 *** |
|  | Diameter | 0.0061 | 7.55E-01 NS |
|  | Global transitivity | -0.1532 | 3.06E-17 *** |
| PPI | Degree | -0.0249 | 8.20E-03 ** |
|  | Hub score | -0.0942 | < 2.2E-16 *** |
|  | Transitivity | -0.0338 | 6.24E-04 *** |
|  | Betweenness | -0.0140 | 1.31E-01 NS |

Note: All correlations are computed using Kendall's rank correlation test, with p-value codes defined as \*\*\* < 0.001 < \*\* < 0.01 < \* < 0.05 < . < 0.1. NS = non-significant. PPI: protein-protein interactions.

By applying similar measures on the PPI network, we report significant negative correlation between F* and PPI centrality measures (**Figure 4H-K**, **Table 2**). Because the PPI network is not directed, authority scores and hub scores cannot be distinguished. The results obtained with the mouse PPI interaction network are qualitatively similar to the ones obtained by Li et al (2010) on Yeast expression data (Li et al. 2010). In addition, we further report that genes involved in complex interactions (that is, genes which interact with more than one other protein simultaneously) have reduced noise in gene expression (Wilcoxon rank test, p-value = 8.053E-05, **Figure 4L**), corroborating previous findings in Yeast (Fraser et al. 2004). Conversely, genes involved in polymeric interactions, that is, where multiple copies of the encoded protein interact with each other, did not show significantly different noise than other genes (Wilcoxon rank test, p-value = 0.0821, **Figure 4M**).

It was previously shown that centrality measures negatively correlate with evolutionary rate (Hahn and Kern 2004). Our results suggest that central genes are selectively constrained for their transcriptional noise, and that centrality therefore also influences the regulation of gene expression. Interestingly, it has been reported that central genes tend to be more duplicated (Vitkup et al. 2006). The authors proposed that such duplication events would have been favored as they would confer greater robustness to deleterious mutations in proteins. Our results are compatible with another, non exclusive, possible advantage: having more gene copies could reduce transcriptional noise by averaging the amount of transcripts produced by each gene copy (Raser and O’Shea 2005).

### Network structure impacts transcriptional noise of constitutive genes

Whereas estimators of node centrality highlight gene-specific properties inside a given network, measures at the whole-network level enable the comparison of networks with distinct properties. We computed the size, diameter and global transitivity for each annotated network in our data set (1,364 networks, Supplementary Material) which we compare with the average F* measure of all constitutive nodes. The size of a network is defined as its total number of nodes, while diameter is the length of the shortest path between the two most distant nodes. Transitivity is a measure of connectivity, defined as the average of all nodes’ clustering coefficients. Interestingly, while network size is positively correlated with average degree and transitivity (Kendall’s tau = 0.5880, p-value < 2.2e-16 and Kendall’s tau = 0.1166, p-value = 1.08E-10, respectively), diameter displays a positive correlation with average degree (Kendall’s tau = 0.2959, p-value < 2.2e-16) but a negative correlation with transitivity (Kendall’s tau = -0.0840, p-value = 2.17E-05). This is because diameter increases logarithmically with size, that is, addition of new nodes to large networks do not increase the diameter as much as additions to small networks. This suggests that larger networks are relatively more compact than smaller ones, and their constitutive nodes are therefore more connected. We find that average transcriptional noise correlates negatively with network size (Kendall’s tau = -0.0514, p-value = 0.0039), while being independent of the diameter (Kendall’s tau = 0.0061, p-value = 0.7547 see **Table 3)**. These results are in line with the node-based analyses, and show that the more connections a network has, the less stochastic the expression of the underlying genes is. This supports the view of Raser and Oshea (Raser and O’Shea 2005) that the gene-extrinsic, pathway-intrinsic level is functionally pertinent and needs to be distinguished from the globally extrinsic level. We further asked whether genes with similar transcriptional noise tend to synthesize proteins that connect to each other (positive assortativity) in a given network, or on the contrary, tend to avoid each other (negative assortativity). We considered all Reactome pathways annotated to the mouse and estimated their respective F* assortativity. We found the mean assortativity to be significantly negative, with a value of -0.1384 (one sample Wilcoxon rank test, p-value < 2.2e-16), meaning that proteins with different F* values tend to connect with each other (**Figure S3**). Maslov & Sneppen (Maslov and Sneppen 2002) reported a negative assortativity between hubs in protein-protein interaction networks, which they hypothesized to be the result of selection for reduced vulnerability to deleterious perturbations. In our data set, however, we find the assortativity of hub scores to be significantly positive (average of 0.1221, one sample Wilcoxon rank test, p-value = 1.212E-12, **Figure S5**), although with a large distribution of assortativity values. As we showed that hub scores correlates negatively with F* (**Table 2**), we asked whether the negative assortativity of hub proteins can at least partly explain the negative assortativity of F*. We found a significantly positive correlation between the two assortativity measures (Kendall’s tau = 0.2581, p-value < 2.2e-16). The relationship between the measures, however, is not linear (**Figure S5**), suggesting a distinct relationship between hub score and F* for negative and positive hub score assortativity. Negative assortativity of hub proteins contributes to a negative assortativity of SGE (Kendall’s tau = 0.2730, p-value < 2.2e-16), while for pathways with positive hub score assortativity the effect vanishes (Kendall’s tau = 0.0940, p-value = 3.135E-4). While assortativity of F* is closer to 0 for pathways with positive assortativity of hub score, we note that it is still significantly negative (average = -0.0818, one sample Wilcoxon test with p-value < 2.2e-16). This suggests the existence of additional constraints that act on the distribution of noisy proteins in a network.

**Table 3:** Linear models of transcriptional noise with genomic and epigenomic factors.

|  | OLS |  |  | GLS |  |  |
| --- | --- | --- | --- | --- | --- | --- |
|  | Coefficient | SE | p-value | Coefficient | SE | p-value |
| (Intercept) | 0.1612 | 0.0781 | 0.0392 * | 0.1665 | 0.0663 | 0.0121 * |
| PC1 | 0.0390 | 0.0065 | <0.0001 *** | 0.0396 | 0.0065 | 0.0000 *** |
| PC2 | -0.0048 | 0.0069 | 0.4854 | -0.0048 | 0.0069 | 0.4838 |
| PC3 | -0.0526 | 0.0091 | <0.0001 *** | -0.0518 | 0.0092 | 0.0000 *** |
| PC4 | -0.0102 | 0.0097 | 0.2905 | -0.0109 | 0.0100 | 0.2773 |
| PC5 | 0.0117 | 0.0106 | 0.2713 | 0.0123 | 0.0106 | 0.2456 |
| PC6 | -0.0152 | 0.0107 | 0.1536 | -0.0152 | 0.0109 | 0.1623 |
| PC7 | 0.0210 | 0.0102 | 0.0384 * | 0.0211 | 0.0110 | 0.0561 . |
| PC8 | 0.0100 | 0.0113 | 0.3778 | 0.0073 | 0.0114 | 0.5250 |
| TFPC1 | 0.0028 | 0.0041 | 0.4912 | 0.0025 | 0.0034 | 0.4658 |
| TFPC2 | 0.0025 | 0.0027 | 0.3664 | 0.0024 | 0.0026 | 0.3585 |
| TFPC3 | 0.0032 | 0.0042 | 0.4513 | 0.0032 | 0.0037 | 0.3825 |
| HistPC1 | -0.0031 | 0.001 | 0.0015 ** | -0.0033 | 0.0010 | 0.0007 *** |
| HistPC2 | -0.0027 | 0.0016 | 0.0846 . | -0.0029 | 0.0015 | 0.0566 . |

Note: OLS: Ordinary Least Squares. GLS: Generalized Least Squares. SE: standard error. Pathway PC1-8: principal components on centrality measures, protein conservation and gene age. TFPC1-3: principal components of the logistic PCA on transcription factors binding evidences. HistPC1 and 2: principal components of the logistic PCA on histone modification marks.

### Transcriptional noise is positively correlated with the evolutionary rate of proteins s

In the yeast *Saccharomyces cerevisiae*, evolutionary divergence between orthologous coding sequences correlates negatively with fitness effect on knock-out strains of the corresponding genes (Hirsh and Fraser 2001), demonstrating that protein functional importance is reflected in the strength of purifying selection acting on it. Fraser et al (Fraser et al. 2004) studied transcription and translation rates of yeast genes and classified genes in distinct noise categories according to their expression strategies. They reported that genes with high fitness effect display lower expression noise than the rest. Following these pioneering observations, we hypothesized that genes under strong purifying selection at the protein sequence level should also be highly constrained for their expression and therefore display a lower transcriptional noise. To test this hypothesis, we correlated F* with the ratio of non-synonymous (Ka) to synonymous substitutions (Ks), as measured by sequence comparison between mouse genes and their human orthologs, after discarding genes with evidence for positive selection (n = 5). In agreement with our prediction, we report a significantly positive correlation between the Ka / Ks ratio and F* (**Figure 4N**, Kendall’s tau = 0.0557, p-value < 1.143E-05), that is, highly constrained genes (low Ka / Ks ratio) display less transcriptional noise (low F*) than fast evolving ones. This result demonstrates that genes encoding proteins under strong purifying selection are also more constrained on their transcriptional noise.

### Older genes are less noisy

Evolution of new genes was long thought to occur via duplication and modification of existing, genetic material (“evolutionary tinkering”, (Jacob 1977)). Evidence for *de novo* gene emergence is however becoming more and more common (Tautz and Domazet-Lošo 2011; Xie et al. 2012). *De novo* created genes undergo several optimization steps, including their integration into a egulatory network (Neme and Tautz 2013). We tested whether the historical process of incorporation of new genes into pathways impacts the evolution of transcriptional noise. We used the phylostratigraphic approach of Neme & Tautz (Neme and Tautz 2013), which categorizes genes into 20 strata, to compute gene age and tested for a correlation with F*. As older genes tend to be more conserved (Wolf et al. 2009), more central (according to the preferential attachment model of network growth (Jeong et al. 2000; Jeong et al. 2001)) and more pleiotropic, we controlled for these confounding factors (Kendall’s tau = -0.0663, p-value = 1.58E-37; partial correlation controlling for Ka / Ks ratio, centrality measures and pleiotropy level, **Figure 4O**). These results suggest that older genes are more deterministically expressed while younger genes are more noisy. While we cannot rule out that functional constraints not fully accounted for by the Ka / Ks ratio or unavailable functional annotations could explain at least partially the correlation of gene age and transcriptional noise, we hypothesise that the observed correlation results from ancient genes having acquired more complex regulation schemes through time. Such schemes include for instance negative feedback loops, which have been shown to stabilize gene expression and reduce expression noise (Becskei and Serrano 2000; Thattai and Oudenaarden 2001).

### Position in the protein network is the main driver of transcriptional noise

In order to jointly assess the effect of network topology, epigenomic factors, Ka / Ks ratio and gene age, we modeled the patterns of transcriptional noise as a function of multiple predictive factors within the linear model framework. This analysis could be performed on a set of 2,794 genes for which values were available jointly for all variables. In order to avoid colinearity issues because some of these variables are intrinsically correlated, we performed data reduction procedures prior to modeling. For continuous variables, including Pathway and PPI network variables, Ka / Ks ratio and gene age, we conducted a principal component analysis (PCA) and used as synthetic measures the first eight principal components (PC), explaining together more than 80% of the total inertia (**Figure S2A**). The first principal component (PC1) of the PCA analysis is associated with pathway centrality measures (degree, hub score, authority score and transitivity, **Figure S2B**). The second principal component (PC2) corresponds to PPI centrality measures (degree, hub score and betweenness), while the third component (PC3) relates to gene age and Ka / Ks ratio. The fourth component (PC4) is associated with PPI complex interactions and transitivity. PC5 and PC6 are essentially associated to betweenness and closeness of the pathway network, PC7 with PPI polymeric interactions and PC8 with pathway pleiotropy. As transcription factors and histone marks data are binary (presence / absence for each gene), we performed a logistic PCA for both type of variables (Landgraf and Lee 2015). For transcription factors, we selected the three first components (hereby noted TFPC), which explained 78% of deviance (**Figure S3A**). The loads on the first component (TFPC1) are all negative, meaning that TFPC1 captures a global correlation trend and does not discriminate between TFs. Tcfcp2l1 appears to be the TF with the highest correlation to TFPC1. The second component TFPC2 is dominated by TCFC (positive loading) and Oct4 (negative loading), while the third component TFPC3 is dominated by Esrrb (positive loading) and MYC, nMyc and E2F1 (negative loadings, **Figure S3B**). For histone marks, the two first components (hereby noted HistPC) explained 95% of variance and were therefore retained (**Figure S4A**). HistPC1 is dominated by marks H3K27me3 linked to gene repression (negative loadings) and HistPC2 by marks H3K4me1 and H3K4me3 linked to gene activation (positive loadings, **Figure S4A**).

We fitted a linear model with F* as a response variable and all 13 synthetic variables as explanatory variables. We find that PC1 has a significant positive effect on F* (**Table 3**). As the loadings of the centrality measures on PC1 are negative (**Figure S2C**), this result is consistent with our finding of a negative correlation of pathway-based centrality measure with F*. PC3 has a highly significant negative effect on F*, which is consistent with a negative correlation with gene age (positive loading on PC3) and a positive correlation with the Ka / Ks ratio (negative loading on PC3, **Figure S2D**). The last highly significant variable is the first principal component of the logistic PCA on histone methylation patterns, HistPC1, which has a negative effect on F*. Because the loadings are essentially negative on HistPC1, this suggests a positive effect of methylation, in particular the repressive H3K27me3. Altogether, the linear model with all variables explained 4.01% of the total variance (adjusted R^2). This small value indicates either that gene idiosyncrasies largely predominate over general effects, or that our estimates of transcriptional noise have a large measurement error, or both. To compare the individual effects of each explanatory variable, we conducted a relative importance analysis. As a mean of comparison, we fitted a similar model with mean expression as a response variable. We find that pathway centrality measures (PC1 variables) account for 38% of the explained variance, while protein constraints and gene age (PC3) account for 32%. Chromatin state (HistPC1) account for another 15% of the variance (**Figure 5**). These results contrast with the model of mean expression, where HistPC1 and HistPC2 respectively account for 51% and 9% of the explained variance, and PC1 and PC3 20% and 10% only (**Figure 5**). This suggests (1) that among all factors tested, position in protein network is the main driver of the evolution of gene-specific stochastic expression, followed by protein constraints and gene age and (2) that different selective pressures act on the mean and cell-to-cell variability of gene expression.

**Figure 5:**
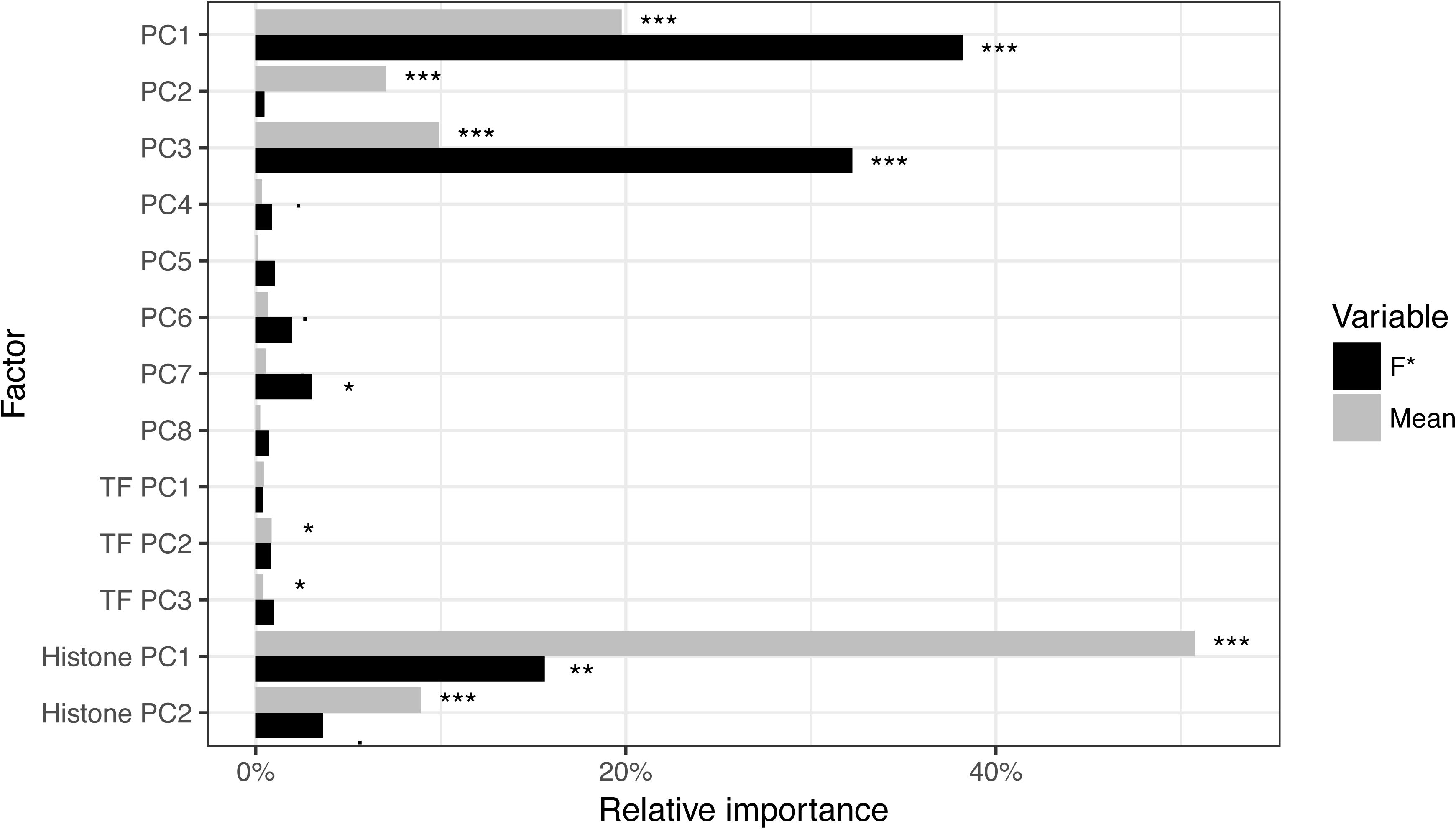
Relative importance of explanatory factors on mean gene expression and F*. Significance codes refer to ANOVA test of variance, *** < 0.001 < ** < 0.01 < * < 0.05 <. < 0.1.

We further included the effect of three-dimensional organization of the genome in order to assess whether it could act as a confounding factor. We developed a correlation model allowing for genes in contact to have correlated values of transcriptional noise. The correlation model was fitted together with the previous linear model in the generalized least square (GLS) framework. This model allows for one additional parameter, λ, which captures the strength of correlation due to three-dimensional organization of the genome (see Methods). The estimate of λ was found to be 0.0016, which means that the spatial autocorrelation of transcriptional noise is low on average. This estimate is significantly higher than zero, and model comparison using Akaike’s information criterion favors the linear model with three-dimensional correlation (AIC = 4880.858 vs. AIC = 4890.396 for a linear model without three-dimensional correlation). Despite the significant effect of 3D genome correlation, our results were qualitatively and quantitatively very similar to the model ignoring 3D correlation (**Table 3**).

### Analysis of bone marrow-derived dendritic cells supports the generality of the results

We assessed the reproducibility of our results by analyzing an additional single-cell transcriptomics data set of 95 unstimulated bone marrow-derived dendritic cells (BMDC) (Shalek et al. 2014). After filtering (see Methods), the data set consisted of 11,640 genes. Using the same normalization procedure as for the ESC data set, we nonetheless report a weak but significant negative correlation between F* and the mean expression, even with a degree-5 polynomial regression (-0.0459, p-value < 1.13E-13). This effect is due to the distribution of per-gene, between cell RFKM values being extremely skewed in this data set. In order to assess the impact of the residual correlation with the mean, we computed a value of F* (noted F_R_*) on a restricted dataset where the variance was between 1/8 and 8 times the mean (75% of all genes) using a quantile regression on the median instead of a linear regression. A second degree polynomial quantile regression proved to be sufficient to remove the effect of mean expression (Kendall’s tau = 0.0114, p-value = 0.1125) on this restricted data set. As all results were consistent when using the F_R_* and F* measures, we only discuss here results obtained with F* and refer to **Supplementary Data 1** for detailed results obtained with the F_R_* measure.

We report a highly significant positive correlation between F* values measured on the 8,792 genes with expression in both data sets, suggesting that cell-to-cell variance in gene expression is to a large extent conserved among the two cell types (Kendall’s tau = 0.1289, p-value < 2.2E-16, **Figure S6A**). GO terms or reactome pathways enrichment analyses reveal less significant but consistant terms with the ESM analysis: the high F* gene set did not show any significantly enriched GO term or reactome pathway (FDR set to 1%) and the low F* gene set revealed RNA-binding as a significantly enriched molecular function, as well as 21 enriched pathways (**Figure S7**). In agreement with results from the ESM analysis, many of the most significant enriched pathways relate to gene expression, including translation and splicing. Interestingly, the two most significant pathways, however, are “Vesicle-mediated transport” and “Membrane trafficking”, two essential pathways for the functioning of dendritic cells. Analyses of network centrality measures also generally show consistent results with the ESC data set, more central genes displaying reduced gene expression noise (**Figure S6B-N, Table S1**). Quantitative differences consists of PPI betweenness, as well as pathway closeness and betweenness are highly significantly negatively correlated with F* while they were only weakly or non-significant with the ESC data set. The only discrepancies that we report between the two data sets relate to pathway level statistics. Pathway size appears to be significantly positively correlated with mean F*, while it was negatively correlated on the ESC data set, yet with a comparatively higher p-value. Similarly pathway diameter is significantly positively correlated with mean F* in the BMDC data set, while it was not significant with the ESC data. We currently have no hypothesis to explain this particular discrepancy. While these results support the generality of our observations, they also illustrate that in details, the fine structure of translational noise may vary in a cell type-specific manner.

We fitted linear models as for the embryonic stem cell (ESC) data set, with the exception that no epigenomic data was available for this cell type. Data reduction was performed using a principal component analysis, with the eight first principal components explaining 81% of the total deviance (**Figure S8A**). We report consistent results with the ESC analysis, with all major effects similar in direction and intensity, highlighting the impact of network centrality measures on expression noise (**Table S2**). With the BMDC data, however, the second principal component PC2 which is associated with PPI centrality measures (**Figure S8B**) appears to have a significant negative impact on F*, while it was not significant with the ESC dataset. As the loading of the PPI centrality measures are positive on PC2, this is consistent with central genes having a lower transcriptional noise as for the pathway network metrics (**Figure S8C**). When taking 3D genome correlations into account, we estimated a low correlation coefficient as for the ESC dataset (lambda = 0.0004), and the AIC favored the model without correlation in this case. Relative importance analysis revealed that network centrality measures contributed most to the explained variance (48% and 21% for PC1 and PC2 respectively), while the contribution of protein constraints and gene age (PC3) was 24%.

### Biological, not technical noise is responsible for the observed patterns

The variance in gene expression measured from single-cell transcriptomics is a combination of biological and technical variance. While the two sources of variance are a priori independent, gene-specific technical variance has been observed in micro-array experiments (Pozhitkov et al. 2007) making a correlation of the two types of variance plausible. If similar effects also affect RNA-Seq experiments, technical variance could be correlated to gene function and therefore act as a covariate in our analyses. In order to assess whether this is the case, we used the dataset of Shalek et al (Shalek et al. 2013), which contains both single-cell transcriptomics and 3 replicates of 10,000 pooled-cell RNA sequencing. In traditional RNA sequencing, which is typically performed on pooled populations of several thousands of cells, biological variance is averaged out so that the resulting measured variance between replicates is essentially the result of technical noise. We computed the mean and variance in expression of each gene across the three populations of cells. By plotting the variance versus the mean in log-space, we were able to compute a “technical” F* 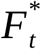) value for each gene (see Methods). We fitted linear models as for the single cell data, using 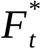 instead of F*. We report that no variable had a significant effect on 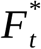 (**Table S3**). In addition, there was no enrichment of the lower 10 ^th^ 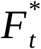 percentile for any particular pathway or GO term. The upper 90^th^ percentile showed no GO term enrichment, but four pathways appeared to be significant: “Chromosome maintenance” (adjusted p-value = 0.0043), “Polymerase switching on the C-strand of the telomere” (adjusted p-value = 0.0062), “Polymerase switching” (adjusted p-value = 0.0062) and “Leading strand synthesis” (adjusted p-value = 0.0062), which relate to DNA replication. While it is unclear why genes involved in these pathways would display higher technical variance in RNA sequencing, these results strikingly differ from our analyses of single cell RNA sequencing and therefore suggest that technical variance does not act as a confounding factor in our analyses.

Because only three replicates were available in the pooled RNA-Seq data set, we asked whether the resulting estimate of mean and variance in expression is accurate enough to allow proper inference of noise and its correlation with other variables. We conducted a jackknife procedure where we sampled the original cells from the ESC data set and re-estimated F* for each sample. We tested combinations of 3, 5, 10 and 15 cells, with 1,000 samples in each case. In each sample, we computed F* with the same procedure as for the complete data set, and fitted a linear model with all 13 synthetic variables. For computational efficiency, we did not include 3D correlation in this analysis. We compute for each variable the number of samples where the effect is significant at the 5% level and has the same sign as in the model fitted on the full data set. We find that the model coefficients are very robust to the number of cells used (**Figure S9A**) and that 3 cells are enough to infer the effect of the PC1 and PC3 variables, the most significant in our analyses. Two main conclusions can be drawn from this jackknife analysis: (1) that the lack of significant effect of our explanatory variables on technical noise is not due to the low number of replicates used to compute the mean and variance in expression and (2) that our conclusions are very robust to the actual cells used in the analysis, ruling out drop-out and amplification biases as possible source of errors (Kharchenko et al. 2014).

### Discussion

Throughout this work, we provided the first genome-wide evolutionary and systemic study of transcriptional noise, using mouse cells as a model. We have shown that transcriptional noise correlates with functional constraints both at the level of the gene itself via the protein it encodes, but also at the level of the pathway(s) the gene belongs to. We further discuss here potential confounding factors in our analyses and argue that our results are compatible with selection acting to reduce noise-propagation at the network level.

In this study, we exhibited several factors explaining the variation in transcriptional noise between genes. While highly significant, the effects we report are of small size, and a complex model accounting for all tested sources of variation only explains a few percent of the total observed variance. There are several possible explanations for this reduced explanatory power: (1) transcriptional noise is a proxy for noise in gene expression, at which selection occurs (**Figure 1**). As transcriptional noise is not randomly distributed across the genome, it must constitute a significant component of expression noise, in agreement with previous observations (Blake et al. 2003; Newman et al. 2006). Translational noise, however, might constitute an important part of the expression noise and was not assessed in this study. (2) Gene expression levels were assessed on embryonic stem cells in culture. Such an experimental system may result in gene expression that differs from that in natural conditions under which natural selection acted. (3) Functional annotations, in particular pathways and gene interactions are incomplete, and network-based measures have most likely large error rates. (4) While the newly introduced F* measure allowed us to assess the distribution of transcriptional noise independently of the average mean expression, it does not capture the full complexity of SGE. Explicit modeling, for instance based in the Beta-Poisson model (Vu et al. 2016) is a promising avenue for the development of more sophisticated quantitative measures.

In a pioneering study, Fraser et al (Fraser et al. 2004), followed by Shalek et al (Shalek et al. 2013), demonstrated that essential genes whose deletion is deleterious, and genes encoding subunits of molecular complexes as well as housekeeping genes display reduced gene expression noise. Our findings go beyond these early observations by providing a statistical assessment of the joint effect of multiple explanatory factors. Our analyses reveal that network centrality measures are the explanatory factors that explained the most significant part of the distribution of transcriptional noise in the genome. Network-based statistics were first tested by Li et al (Li et al. 2010) using PPI data in Yeast. While we are able to extend these results to mouse cells, we show that more detailed annotation as provided by the Reactome database lead to new insights into the selective forces acting on expression noise. Our results suggest that “pathways” constitute a relevant systemic level of organisation, at which selection can act and drive the evolution of SGE at the gene level. This multi-level selection mechanism, we propose, can be explained by selection against noise propagation within networks. It has been experimentally demonstrated that expression noise can be transmitted from one gene to another gene with which it is interacting (Pedraza and van Oudenaarden 2005). Large noise at the network level is deleterious (Barkai and Leibler 1999) but each gene does not contribute equally to it, thus the strength of selective pressure against noise varies among genes in a given network. We have shown that highly connected, “central” proteins typically display reduced transcriptional noise. Such nodes are likely to constitute key players in the flow of noise in intra-cellular networks as they are more likely to transmit noise to other components. In accordance with this hypothesis, we find genes with the lowest amount of transcriptional noise to be enriched for top-level functions, in particular involved in the regulation of other genes.

These results have several implications for the evolution of gene networks. First, this means that new connections in a network can potentially be deleterious if they link genes with highly stochastic expression. Second, distinct selective pressures at the “regulome” and “interactome” levels (**Figure 1**) might act in opposite direction. We expect genes encoding highly connected proteins to have more complex regulation schemes, in particular if their proteins are involved in several biological pathways. In accordance, several studies demonstrated that expression noise of a gene positively correlates with the number of transcription factors controlling its regulation (Sharon et al. 2014), a correlation that we also find significant in the data set analyzed in this work. Central genes, while being under negative selection against stochastic behavior, are then more likely to be controlled by numerous transcription factors which increase transcriptional noise. As a consequence, if the number of connections at the interactome level is correlated with the number of connections at the regulome level, we predict the existence of a trade-off in the number of connections a gene can make in a network. Alternatively, highly connected genes might evolve regulatory mechanisms allowing them to uncouple these two levels: negative feedback loops, for instance, where the product of a gene down-regulates its own production have been shown to stabilize expression and significantly reduce stochasticity (Becskei and Serrano 2000; Dublanche et al. 2006; Tao et al. 2007). We therefore predict that negative feedback loops are more likely to occur at genes that are more central in protein networks, as they will confer greater resilience against high SGE, which is advantageous for this class of genes.

Our results enabled the identification of possible selective pressures acting on the level of stochasticity in gene expression. The mechanisms by which the amount of stochasticity can be controlled remain however to be elucidated. We evoked the existence of negative feedback loops which reduce stochasticity and the multiplicity of upstream regulator which increase it. Recent work by Wolf et al (Wolf et al. 2015) and Metzger et al (Metzger et al. 2015) add further perspective to this scheme. Wolf and colleagues found that in *Escherichia coli* noise is higher for natural than experimentally evolved promoters selected for their mean expression level. They hypothesized that higher noise is selectively advantageous in case of changing environments. On the other hand, Metzger and colleagues performed mutagenesis experiments and found signature of selection for reduced noise in natural populations of *Saccharomyces cerevisae.* These seemingly opposing results combined with our observations provide additional evidence that the amount of stochasticity in the expression of single genes has an optimum, as high values are deleterious because of noise propagation in the network, whilst lower values, which result in reduced phenotypic plasticity, are suboptimal in case of dynamic environment.

### Conclusion

Using a new measure of transcriptional noise, our results demonstrate that the position of the protein in the interactome is a major driver of selection against stochastic gene expression. As such, transcriptional noise is an essential component of the phenotype, in addition to the mean expression level and the actual sequence and structure of the encoded proteins. This is currently an under-appreciated phenomenon, and gene expression studies that focus only on the mean expression of genes may be missing key information about expression diversity. The study of gene expression must consider changes in noise in addition to change in mean expression level as a putative explanation for adaptation. Further work aiming to unravel the exact structure of the regulome is however needed in order to fully understand how transcriptional noise is generated or inhibited.

## Material and Methods

### Single-cell gene expression data set

We used the dataset generated by Sasagawa et al. (Sasagawa et al. 2013) retrieved from the Gene Expression Omnibus repository (accession number GSE42268). We analyzed expression data corresponding to embryonic stem cells in G1 phase, for which more individual cells were sequenced. A total of 17,063 genes had non-zero expression in at least one of the 20 single cells. Similar to Shalek et al (Shalek et al. 2014), a filtering procedure was performed where only genes whose expression level satisfied log(FPKM+1) > 1.5 in at least one single cell were kept for further analyses. This filtering step resulted in a total of 13,660 appreciably expressed genes for which transcriptional noise was evaluated.

### Measure of transcriptional noise

The expression mean (μ) and variance (*σ* ^2^) of each gene over all single cells were computed. We measured stochastic gene expression as thee ratio 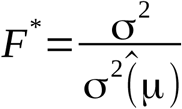, where 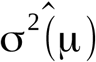 is the expected variance given the mean expression. In order to compute 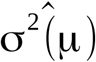, we performed several polynomial regressions with log(*σ* ^2^) as a function of log(μ), with degrees between 1 and 5. We then tested the resulting F* measures for residual correlation with mean expression using Kendall’s rank correlation test. We find that a degree-3 polynomial regression was sufficient to remove any residual correlation with F* (Kendall’s tau = 0.0037, p-value = 0.5217). F* can be seen as a general expression for the Fano factor and noise measure: when using a polynome of degree 1, the expression of F* becomes 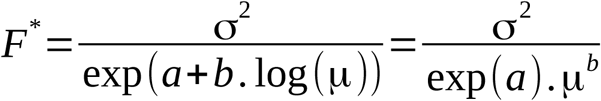, and is therefore equivalent to the Fano factor when a = 0 and b = 1, and equivalent to noise when a = 0 and b = 2.

### Genome architecture

The mouse proteome from Ensembl (genome version: mm9) was used in order to get coordinates of all genes. The Hi-C dataset for embryonic stem cells (ES) from Dixon et al (Dixon et al. 2012) was used to get three-dimensional domain information. Two genes were considered in proximity in one dimension (1D) if they are on the same chromosome and no protein-coding gene was found between them. The primary distance (in number of nucleotides) between their midpoint coordinates was also recorded as 1D a distance measure between the genes. Two genes were considered in proximity in three dimensions (3D) if the normalized contact number between the two windows the genes belong was non-null. Two genes belonging to the same window were considered in proximity. We further computed the relative difference of stochastic gene expression between two genes by computing the ratio 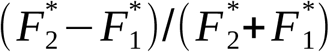. For each chromosome, we independently tested if there was a correlation between the primary distance and the relative difference in stochastic gene expression with a Mantel test, as implemented in the ade4 package (Dray and Dufour 2007). In order to test whether genes in proximity (1D and 3D) had more similar transcriptional noise than distant genes, we contrasted the relative differences in transcription noise between pairs of genes in proximity and pairs of distant genes. As we test all pairs of genes, we performed a randomization procedure in order to assess the significance of the observed differences by permuting the rows and columns in the proximity matrices 10,000 times. Linear models accounting for spatial interactions with genes were fitted using the generalized least squares (GLS) procedure as implemented in the “nlme” package for R. A correlation matrix between all tested genes was defined as *G*={*gi, j*}, where *gi, j* is the correlation between genes i and j. We defined *gi, j*=1-exp(-λ δ*i, j)*, where δ*i, j* takes 1 if genes i and j are in proximity, 0 otherwise (binary model). Alternatively, δ*i, j* can be defined as the actual number of contacts between the two 20 kb regions (as defined by Dixon et al) the genes belong to (proportional model). Parameter l was estimated jointly with other model parameters, it measures the strength of the genome “spatial” correlation. Models were compared using Akaike’s information criterion (AIC). We find that the proportional correlation model fitted the data better and therefore selected it for further analyses.

### Transcription factors and histone marks

Transcription factor (TF) mapping data from the Ensembl regulatory build (Zerbino et al. 2015) were obtained via the biomaRt package for R. We used the Grch37 build as it contained data for stem cells epigenomes. Genes were considered to be associated with a given TF when at least one binding evidence was present in the 3 kb upstream flanking region. Transcription factors associated with less than 5 genes for which transcriptional noise could be computed were not considered further. A similar mapping was performed for histone marks by counting the evidence of histone modification in the 3 kb upstream and downstream regions of each gene. A logistic principal component analysis was conducted on the resulting binary contingency tables using the logisticPCA package for R (Landgraf and Lee 2015), for TF and histone marks separately. Principal components were used to define synthetic variables for further analyses.

### Biological pathways, protein-protein interactions and network topology

We defined genes either in the top 10% least noisy or in the top 10% most noisy as candidate sets and used the Reactome PA package (Yu and He 2016) to search the mouse Reactome database for overrepresented pathways with a 1% false discovery rate.

Centrality measures were computed using a combination of the “igraph” (Csardi and Nepusz 2006) and “graphite” (Sales et al. 2012) packages for R. As the calculation of assortativity does not handle missing data (that is, nodes of the pathway for which no value could be computed), we computed assortativity on the sub-network with nodes for which data were available. Reactome centrality measures could be computed for a total of 4,454 genes with expression data.

Protein-protein interactions (PPI) were retrieved from the iRefIndex database (Razick et al. 2008) using the iRefR package for R (Mora and Donaldson 2011). Interactions were converted to a graph using the dedicated R functions in the package, and the same methods were used to compute centrality measures as for the pathway analysis. Because the PPI-based graph was not oriented, authority scores were not computed for this data (as this gave identical results to hub scores). Furthermore, as most genes are part of a single graph structure in the case of PPI interactions, closeness values were not further analysed as they were virtually identical for all genes.

### Gene Ontology Enrichment

Eight thousand three hundreds and twenty five out of the 13,660 genes were associated with Gene Ontology (GO) terms. We tested genes for GO terms enrichment at both ends of the F* spectrum using the same threshold percentile of 10% low / high noise genes as we did for the Reactome analysis. We carried out GO enrichment analyses using two different algorithms: “Parent-child” (Grossmann et al. 2007) and “Weight01”, a mixture of two algorithms developed by Alexa et al (Alexa et al. 2006). We kept only the terms that appeared simultaneously on both Parent-child and Weight01 under 1% significance level, controlling for multiple testing using the FDR method (Benjamini and Hochberg 1995).

### Sequence divergence

The Ensembl’s Biomart interface was used to retrieve the proportion of non-synonymous (Ka) and synonymous (Ks) divergence estimates for each mouse gene relative to the human ortholog. This information was available for 13,136 genes.

### Gene Age

The relative taxonomic ages of the mouse genes have been computed and is available in the form of 20 Phylostrata (Neme and Tautz 2013). Each Phylostratum corresponds to a node in the phylogenetic tree of life. Phylostratum 1 corresponds to “All cellular organisms” whereas Phylostratum 20 corresponds to “*Mus musculus*”, with other levels in between. We used this published information to assign each of our genes to a specific Phylostratum and used this as a relative measure of gene age: Age = 21 - Phylostratum, so that an age of 1 corresponds to genes specific to *M. musculus* and genes with an age of 20 are found in all cellular organisms.

### Linear modeling

We simultaneously assessed the effect of different factors on transcriptional noise by fitting linear models to the gene-specific F* estimates. To avoid colinearity issues of intrinsically correlated explanatory variables, we conducted a data reduction procedure using multivariate analysis. We used variants of principal component analysis (PCA) on explanatory variables in three groups: network centrality measures, Ka / Ks and gene age with standard PCA, transcription factor binding evidence and histone methylation patterns using logistic PCA, a generalization of PCA for binary variables (Landgraf and Lee 2015). In each case, we used the most representative components (totaling at least 75% of the total deviance) as synthetic variables. PCA analysis was conducted using the ade4 package for R (Dray and Dufour 2007), logistic PCA was performed using the logisticPCA package (Landgraf and Lee 2015).

We built a linear model with F* as a response variable and thirteen synthetic variables as explanatory variables. As the synthetic variables are principal components, they are orthogonal by construction. The fitted model displayed significant departure to normality and was further transformed using the Box-Cox procedure (“boxcox” function from the MASS package for R (Venables and Ripley 2002)). Residues of the selected model had normal, independent residue distributions (Shapiro-Wilk test of normality, p-value = 0.121, Ljung-Box test of independence, p-value = 0.2061) but still displayed significant heteroscedasticity (Harrison-McCabe test, p-value = 0.003). In order to ensure that this departure from the Gauss-Markov assumptions does not bias our inference, we used the “robcov” function of the “rms” package in order to get robust estimates of the effect significativity (Harrell 2015). Relative importance of each explanatory factor was assessed using the method of Lindeman, Merenda and Gold (Lindeman et al. 1979) as implemented is the R package “relaimpo”. The significance of the level of variance explained by each factor was computed using standard ANOVA procedure.

### Additional data sets

The aforementioned analyses were additionally conducted on the bone marrow-derived dendritic cells data set of Shalek et al (Shalek et al. 2014). Following the filtering procedure established by the authors in the original paper, genes which did not satisfied the condition of being expressed by an amount such that log(TPM+1) > 1 in at least one of the 95 single cells were further discarded, where TPM stands for transcripts per million. This cut-off threshold resulted in 11,640 genes being kept for investigation. The rest of the analyses was conducted in the same way as for the ESM data set.

### Jackknife procedure

A jackknife procedure was conducted in order to assess (1) the robustness of our results to the choice of actual cells used to estimate mean and variance in gene expression and (2) the power of the pooled RNA-seq analysis for which only three replicates were available. This analysis was conducted by sampling 3, 5, 10 and 15 of the original 20 single cells of the ESM data set (Sasagawa et al. 2013), 1,000 times in each case. The exact same analysis was conducted on each random sample as for the complete data set, and model coefficients and their associated p-values were recorded.

### Data and program availability

All datasets and scripts to reproduce the results of this study are available under the DOI 10.6084/m9.figshare.4587169.

### Authors contributions

GVB and JYD designed the experiments and wrote the manuscript. GVB, NP and JYD conducted the analyses.

## Acknowledgements

The authors would like to thank Rafiq Neme-Garrido, Frederic Bartels and Estelle Renaud for fruitful discussions about this work, Andrew Landgraf for help with the logistic PCA analysis as well as Diethard Tautz for comments on an earlier version of this manuscript. JYD acknowledges funding from the Max Planck Society. This work was supported by the German Research Foundation (DFG), within the priority program (SPP) 1590.

## Supplementary material

**Table S1**: Correlation of transcriptional noise with genes centrality measures and pleiotropy for the bone marrow-derived dendritic cells data set. Legends as in **Table 2**.

**Table S2**: Linear models of transcriptional noise with genomic factors for the bone marrow-derived dendritic cells data set. Legend as in Table 4.

**Table S3**: Linear model of transcriptional noise with genomic factors with pooled RNA-Seq data. Legend as in Table 4.

**Figure S1**: Impact of genome organization on the distribution of transcriptional noise. The x-axis shows the mean relative difference in transcriptional noise. Vertical lines show observed values and histograms the distribution over 10,000 permutations (see Methods). Left panel: distribution for neighbor genes along the genome. Right panel: distribution for genes in contact in three-dimensions.

**Figure S2**: Principal component analysis of pathways centrality measures. A) Proportion of deviance explained by models with 1, 2, etc. principal components. B) Contributions, computed as proportion of deviance, of each input variable to each principal component. C) Loadings of each variable on the 2 first components. D) Loadings of each variable on the 3rd and 4th principal components.

**Figure S3**: Logistic principal component analysis of transcription factor binding evidences. A) Proportion of deviance explained by models with 1, 2, etc. principal components. B) Contributions, computed as proportion of deviance, of each input variable to each principal component. C) Loadings of each variable on the 2 first components. D) Loadings of each variable on the 2nd and 3rd principal components.

**Figure S4**: Logistic principal component analysis of histone marks. A) Proportion of deviance explained by models with 1, 2, etc. principal components. B) Contributions, computed as proportion of deviance, of each input variable to each principal component. C) Loadings of each variable on the 2 first components.

**Figure S5**: Assortativity in networks. A) Distribution of assortativity values for hub scores. B) Distribution of assortativity values for F*. C) Assortativity for F* and hub scores are plotted against each other. Solid lines represent linear regressions fitted on pathways with negative or positive hub score assortativity, respectively. Dashed line represents a linear regression fitted on all data.

**Figure S6**: Factors driving stochastic gene expression in the bone marrow-derived dendritic cells data set. Legends as in **Figure 4**.

**Figure S7**: Enriched pathways in the low-noise gene set of the bone marrow-derived dendritic cells data set.

**Figure S8**: Principal component analysis of pathways centrality measures of the bone marrow-derived dendritic cells data set. Legends as in **Figure S2**.

**Figure S9**: Robustness and power analysis. A jackknife procedure was conducted by fitted linear models with all explanatory variables on a subset of cells taken randomly (x-axis). A) estimated coefficient of each effect. B) proportion of simulations where the coefficient is significant at the 5% level. Filled bars correspond to significant effect when the complete data set is used. PC: principal component. PPI: protein-protein interactions. TF: transcription factors.

**Supplementary Data 1**: All scripts and data set necessary to reproduce the analyses and figures in this manuscript.

